# Autocrine TGFβ2 enforces a transcriptionally hybrid cell state in Ewing sarcoma

**DOI:** 10.1101/2025.06.11.653876

**Authors:** Emma D. Wrenn, Jacob C. Harris, April A. Apfelbaum, Jonah R. Valenti, Patricia A. Lipson, Stephanie I. Walter, Nicolas M. Garcia, Aya Miyaki, Steven C. Chen, Jim M. Olson, Jason S. Price, Kelly M. Bailey, Elizabeth R. Lawlor

**Author notes:** Corresponding authors: Elizabeth R. Lawlor,; Emma D. Wrenn.

## Abstract

Sub-populations of cancer-associated fibroblast (CAF)-like tumor cells deposit extracellular matrix (ECM) proteins that support Ewing sarcoma (EwS) progression and metastasis. We previously showed a hallmark of CAF-like EwS cells is their hybrid transcriptional state wherein the driver fusion oncogene, EWS::FLI1, maintains activation of proliferative programs but loses capacity to repress mesenchymal genes. Here, we studied primary patient tumors and cell line models to identify molecular drivers of this hybrid state. Our data reveal that hybrid EwS cells are induced and maintained by a TGFβ signaling positive feedback loop. Hybrid cells de-repress *TGFBR2* and upregulate expression and secretion of TGFβ2 to sustain pathway activation and ECM deposition. While TGFβ ligands can potently induce growth arrest in cells of epithelial origin, we show that TGFβ1 and TGFβ2 promote cell invasion of EwS cells without affecting proliferation. Thus, stroma and tumor-derived TGFβ ligands induce and maintain hybrid EwS cells to promote pro-metastatic cell phenotypes.

## Introduction

Cell plasticity plays critical roles during cancer progression, giving cancer cells access to distinct phenotypic cell states that allow tumors to adapt and thrive (*1*). This plasticity contributes to major clinical barriers including tumor heterogeneity, acquisition of metastatic potential, and escape from therapy (*2–5*). Recent studies have highlighted the cell state heterogeneity of Ewing sarcoma (EwS), an aggressive bone and soft tissue tumor driven by pathognomonic FET::ETS transcription factor fusions, most commonly EWS::FLI1 (*6, 7*). EwS samples show remarkable tumor cell heterogeneity in expression of mesenchymal identity genes, including subpopulations that are quasi-neuroectodermal and other highly mesenchymal, ECM secreting subpopulations (*8–10*). Importantly, genes upregulated in more mesenchymal EwS cell subpopulations are associated with metastatic potential and treatment resistance (*11–14*). While survival rates for localized EwS have risen with changes to standard-of-care and intensification of therapy, the 5-year survival rate for metastatic EwS remains <30% (*6*). Elucidating the molecular drivers of cell plasticity therefore has the potential to unveil new therapeutic strategies that could be leveraged to prevent or eliminate EwS tumor cells that promote metastasis.

Mesenchymal identity in EwS cells is largely governed by the transcriptional activity of driver EWS::ETS fusions (reviewed in (*15*)). EWS::FLI1 upregulates thousands of target genes directly at de novo enhancers and indirectly through regulation of other transcription factors and signaling pathways (*16–19*). EWS::FLI1 can also directly and indirectly induce gene repression (*15, 17*). Mesenchymal identity genes are overrepresented among EWS::FLI1 repressed targets, and reduction in EWS::FLI1 expression generates EWS::FLI1-low cell states with increased mesenchymal identity and invasive capacity (*13, 14, 20*). Conversely, EWS::FLI1-high states maintain quasi-neural gene expression and high proliferation (*8, 13, 21*). Using single cell sequencing analyses, we recently highlighted a third subpopulation in EwS models, EWS::FLI1 hybrid cells (*10*). In these cells EWS::FLI1 is still highly expressed and maintains activation of its upregulated target genes, but many downregulated targets are concurrently de-repressed. In particular, EWS::FLI1 hybrid cells upregulate ECM remodeling genes and adopt phenotypes similar to those of cancer-associated fibroblasts (CAFs), leading us to term them CAF-like tumor cells. Importantly, we showed that CAF-like EwS cells retain both proliferative and tumorigenic potential coincident with enhanced migration and ECM deposition (*10*). The hybrid nature of CAF-like EwS cells is thus reminiscent of transitional or hybrid epithelial to mesenchymal transition (EMT) cells in carcinoma, which have recently been implicated as drivers of metastasis (*22, 23*).

Our prior work showed that CAF-like EwS cells are plastic and exist as subpopulations of cells that are highly heterogeneous with respect to frequency and spatial distribution, both within and between EwS models and primary tumors (*10*). Notably, in PDX and xenograft models CAF-like cells were enriched at tumor-stromal borders and adjacent to areas of necrosis suggesting that signals from the tumor microenvironment (TME) might direct EwS cells towards the CAF-like state. Hypoxia, Wnt ligands, IL-6, and other exogenous factors all induce phenotypic and transcriptional changes in EwS cells (*24–29*), supporting the hypothesis that crosstalk with the TME may be a key driver of EwS plasticity and cell state. In the current work, we sought to determine if and how signals from the TME induce EwS cells to adopt transcriptionally hybrid, CAF-like states. Our results identify TGFβ signaling as a master regulator of the hybrid cell state in EwS cells. Specifically, we identify a TGFβ activation loop that is initiated in tumor cells that have lost EWS::FLI1-mediated repression of *TGFBR2*. These TGFβ-responsive tumor cells upregulate expression and secretion of TGFβ2, potentiating TGFβ-induced ECM programs and loss of EWS::FLI1-mediated gene repression whilst having no impact on cell proliferation. These findings identify tumor-derived TGFβ2 as a key lever that amplifies TGFβ-induced cell plasticity and governs transition of EwS cells to transcriptionally hybrid CAF-like states.

## Results

### TGFβ induces CAF-like gene expression in EwS cells

To nominate TME signaling programs that might induce plastic EwS cells to switch to more mesenchymal CAF-like states, we interrogated scRNA-seq data from 9 EwS cell lines (*10*) and 5 patient-derived xenografts (PDXs) (*9*) (**Fig S1A-B**). Gene ontology analysis of genes that were reproducibly and significantly upregulated in single CAF-like cells in vitro and in vivo confirmed high expression of matrix remodeling genes and revealed specific enrichment of terms related to TGFβ signaling (**Fig 1A, B**). Given the established role of TGFβ as an activator of myofibroblastic CAFs and epithelial-mesenchymal transitions in carcinoma (*1, 30, 31*), we hypothesized that TGFβ signaling might similarly activate a CAF-like state in EwS. To test this, we selected a panel of 4 EwS cell lines with very low (TC71), heterogeneous (A673, CHLA10) and high (PDX305) CAF-like gene expression by single cell sequencing (**Fig 1C**). We measured expression of *NT5E* (CD73) and *COL1A1* as markers of the CAF-like state and found that both genes were reproducibly induced by TGFβ1 in all cell lines except for TC71, where response was weak (**Fig 1D**). Notably, TC71 cells harbor relatively few CAF-like cells in basal culture (**Fig 1C** and (*10*)), leading us to speculate that CD73^+^, CAF-like cells are more responsive to TGFβ than bulk CD73-cells. To test this, we measured the impact of TGFβ1 on expression of *NT5E* in A673 and CHLA10 cells that had been sorted into CD73^+^ and CD73^-^ populations. As shown (**Fig 1E)**, *NT5E* expression increased in both populations but both baseline and induced expression were lower in CD73^-^ cells. This suggests that CAF-like cell subpopulations are inherently more responsive to TGFβ signals than the dominant CD73^-^ cells. In further support of this, TGFβ1-induced secretion of the ECM protein Tenascin-C (*TNC*) was minimal in TC71 cells but robust in the other cell lines (**Fig 1F**). Despite this heterogeneity in phenotypic responses, all cell lines activated the TGFβ signaling axis in response to ligand exposure, as demonstrated by phospho-SMAD2 (pSMAD2) expression (**Fig 1G**). Moreover, vactosertib, a small molecule inhibitor of TGFBR1 activity (*32*), greatly reduced TNC deposition by TGFβ1-responsive cell lines (**Fig 1F**). Together these results show that TGFβ can induce SMAD activation in EwS cell lines but the capacity to induce transcription of downstream gene targets and CAF-like phenotypes is highly heterogeneous and is correlated with the basal level of CAF-like identity in each model.

**Fig 1.**
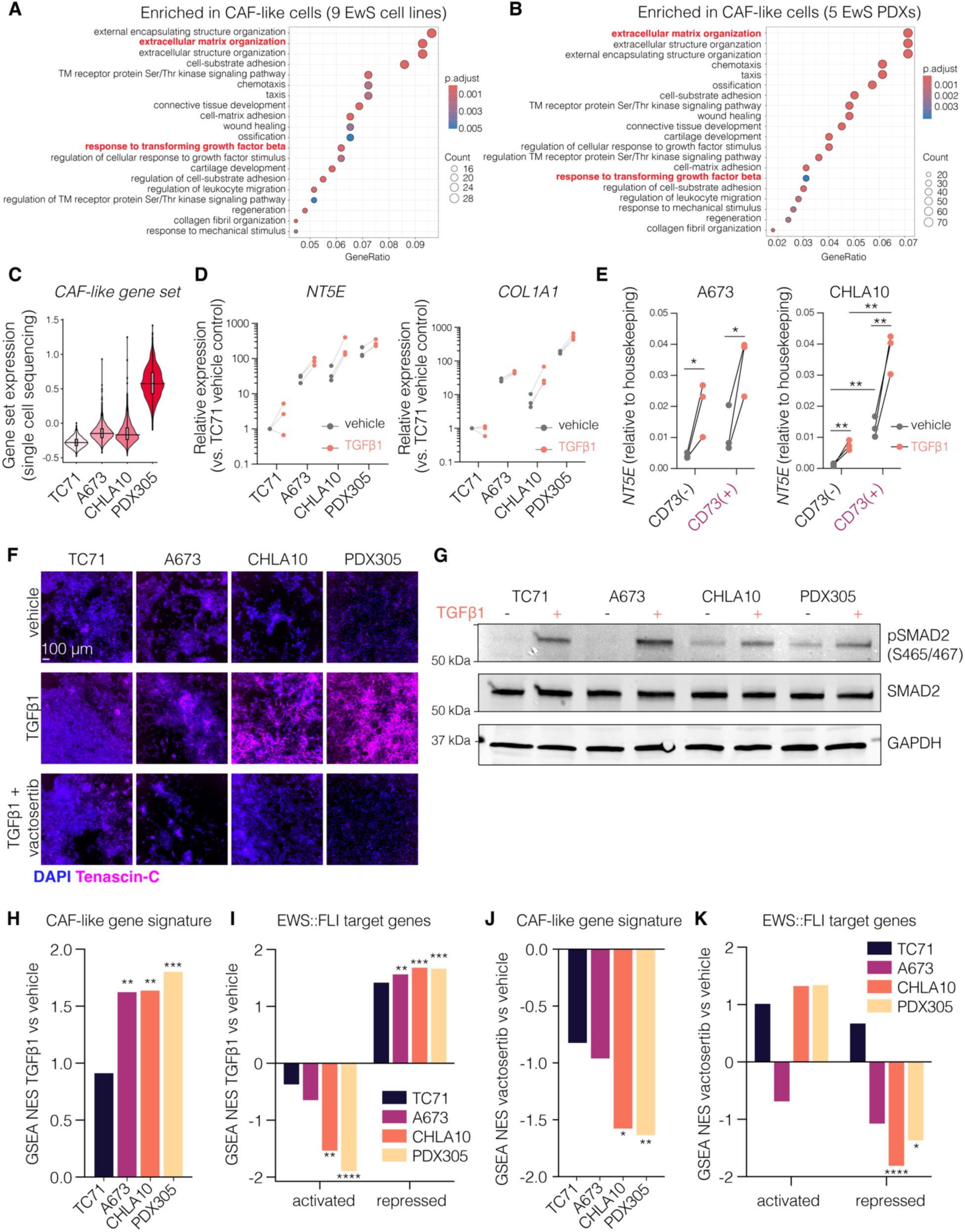
TGFβ induces CAF-like gene expression in EwS cells. Gene ontology of genes marking CAF-like cells (top 30% of cells by expression of 28-gene signature (*10*), marker genes padj <0.05) via single cell sequencing (scSeq) of (**A**) 9 Ewing sarcoma (EwS) cell lines, and (**B**) 5 EwS PDXs in NSG mice (*9*). (**C**) 28-gene EwS CAF-like signature expression in 4 EwS cell lines by scSeq (*10*). (**D**) *TGFBI* and *COL1A1* expression by RT-qPCR after 48 hrs TGFβ1 (10 ng/mL). Normalized to vehicle treated TC71 sample. n=3, lines indicate same biological replicate. (**E**) *NT5E* expression by RT-qPCR in CD73(-) vs. CD73(+) cells treated for 24 hours post-FACS with 10 ng/mL TGFB1. n=3, p-values = unpaired t-tests. (**F**) Immunofluorescence after 72 hrs TGFβ1 (10 ng/mL) -/+ vactosertib (1 μM). Representative of n=2. (**G**) Immunoblots of phospho vs. total SMAD2 in EwS cells treated with TGFB1 (10 ng/mL, 48hrs). Representative of n=2. Loading control = GAPDH. (**H**) GSEA NES scores of 28-gene CAF-like signature in TGFβ1 vs. vehicle treated cells (p-values = padj). (**I**) GSEA of 100 EWS::FLI1 activated or repressed enhancer target genes (*17*) in TGFβ1 vs. vehicle treated cells (p-values = padj). (**J**) GSEA NES scores of 28-gene CAF-like signature in vactosertib vs. vehicle treated cells (p-values = padj). (**K**) GSEA of 100 EWS::FLI1 activated or repressed enhancer target genes (*17*) in vactosertib vs. vehicle treated cells (p-values = padj). ns, *p* > 0.05; *, *p* < 0.05; **, *p* < 0.01; ***, *p* < 0.001; ****, *p* < 0.0001.

To further explore heterogeneity of response in an unbiased fashion, we performed bulk RNA sequencing on 4 cell lines in the presence of TGFβ1 or vactosertib (**Supplemental Table 1**). Consistent with our RT-qPCR findings, TGFβ1 had minimal impact on the TC71 transcriptome but induced robust changes in A673, CHLA10, and PDX305 cell lines (**Fig S1C**). Gene set enrichment analysis (GSEA) confirmed that the 28-gene EwS CAF-like signature (*10*) was induced by TGFβ1 in the 3 responsive models (**Fig 1H**). The transcriptional state of CAF-like EwS cells is characterized by loss of EWS::FLI1-mediated gene repression, with or without concomitant loss of EWS::FLI1-mediated gene activation (*10*). We therefore evaluated expression of published EWS::FLI1 gene target signatures (*17*) in the TGFβ1-responsive cells. Genes that are normally directly repressed by EWS::FLI1 were significantly and reproducibly upregulated by TGFβ1 in A673, CHLA10, and PDX305 cell lines (**Fig 1I**). The EWS::FLI1-activated signature was downregulated by TGFβ1 in CHLA10 and PDX305 cells but not significantly altered in TC71 or A673 cells (**Fig 1I**). Vactosertib treatment significantly downregulated the CAF-like signature (**Fig 1J**) and the EWS::FLI1 repressed signature (**Fig 1K**) in CHLA10 and PDX305 cells, but did not significantly alter the EWS::FLI1 activated signature in any cell line (**Fig 1K**). Thus, these studies show that TGFβ-mediated changes to EwS transcriptomes result in generalized loss of EWS::FLI1-dependent gene repression and a more limited loss of EWS::FLI1-dependent gene activation, consistent with the hybrid transcriptional state of CAF-like cells.

### EwS cells with active TGFβ signaling are detected in cell subpopulations in tumors

Having established that TGFβ1 can induce CAF-like phenotypes and hybrid transcriptional states in vitro, we next performed immunofluorescence staining of a tumor tissue microarray (TMA) to assess TGFβ pathway activation in patient tumors in vivo. CD99 (to mark tumor cell membranes) and pSMAD2 (to mark active TGFβ signaling) were assessed by co-immunofluorescence across 27 patient biopsies (**Supplemental Table 2**). Considerable intra- and inter-tumor heterogeneity of pSMAD2 expression was detected. Though most biopsy cores harbored few dual positive CD99+/pSMAD2+ cells, pSMAD2+ tumor cells were abundant in some samples (**Fig 2A, B**). In 9 of 27 evaluable tumor cores, at least 5% of the CD99+ tumor nuclei were also strongly pSMAD2+ (**Fig 2C**). TNC was most robustly detected in biopsies with higher proportions of pSMAD2+ tumor cells (**Fig 2D**). TNC can be deposited by either tumor cells or stromal cells (*33*) so it is impossible to know if pSMAD2+ EwS cells were the source of TNC in these stroma-rich patient biopsies. As an orthogonal approach to address this question, we stained A673 tumor xenografts with a human-specific anti-TNC antibody and confirmed that TNC adjacent to pSMAD2+ EwS cells was tumor cell-derived (**Fig 2E**). Thus, TGFβ-activated tumor cells can be identified in patient tumors and in EwS xenografts, and these cells are heterogeneous in both abundance and spatial distribution.

**Fig 2.**
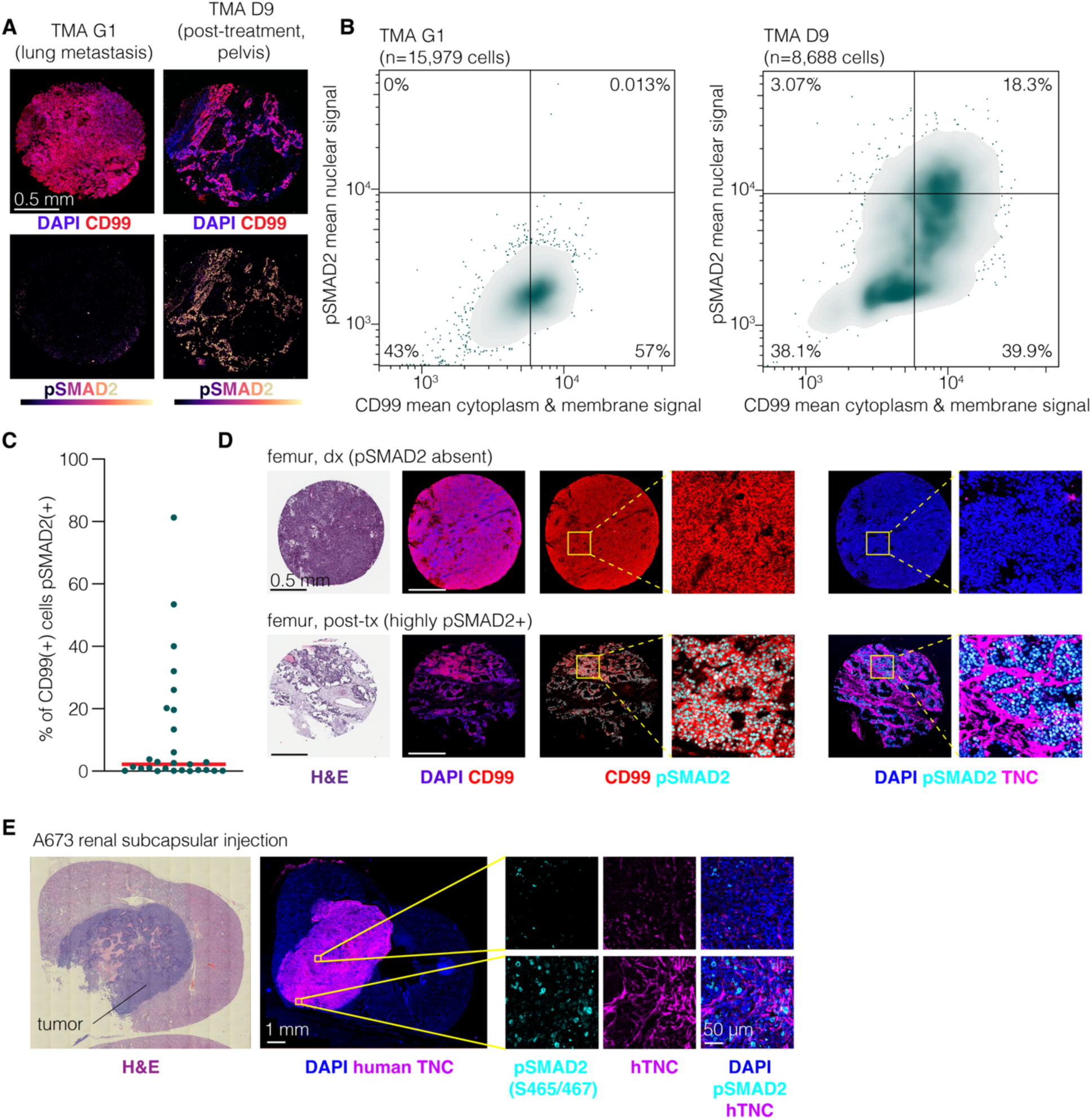
EwS cells with active TGFβ signaling are detected in cell subpopulations in tumors. **(A**) EwS TMA sections stained for CD99 (tumor marker) with low (left) and high (right) pSMAD2 S465/S467 signal. (**B**) Nuclear pSMAD2 S465/467 and cytoplasmic/membranous CD99 average intensity per cell. Gates define positive vs. negative populations (conservative threshold of pSMAD2 or CD99 true positives). (**C**) % CD99+ cells that were pSMAD2+ for each section of the TMA (n=27, Supplemental Table 2), red line = median. (**D**) H&E and IF from EwS TMA sections stained for CD99, pSMAD2 S465/467, and TNC (ECM protein). Right, zoomed insets showing CD99^hi^/pSMAD^low^/TNC^low^ region vs.CD99^hi^/pSMAD2^hi^/TNC^hi^ region. (**E**) H&E of tumor formed 3-weeks post renal subcapsular injection A673 cells in NSG mice. Right, IF for pSMAD2 S465/467 and ECM marker TNC (human specific antibody).

### EWS::FLI1-low and -hybrid transcriptional states escape *TGFBR2* repression

EWS::FLI1 directly represses transcription of *TGFBR2* (*17, 34, 35*). In EWS::FLI1-low cells, *TGFBR2* is derepressed and cells are sensitized to TGFβ signals (*26*). We therefore next sought to determine if *TGFBR2* is de-repressed and a key determinant of the TGFβ response in transcriptionally hybrid EwS cells. In single cell sequencing data of 9 EwS lines we observed minority subpopulations of cells with detectable *TGFBR2* expression ranging from 0.63% (SKNMC) to 30.57% (A673) of cells (**Fig S2A**). To assess *TGFBR2* expression in EWS::FLI1-hybrid relative to EWS::FLI1-low and EWS::FLI1-high (bulk) cells, we informatically scored single cell transcriptomes as high, low, or hybrid, defining EWS::FLI1-hybrid cells as those which expressed both direct EWS::FLI1 activated targets and direct EWS::FLI1 repressed targets (*17*) at levels above the median **(Fig S2B**). Hybrid cells were identified using 6 cell lines, ranging in frequency from 3.04% to 33.8 % (**Fig 3A, Fig S2C**). *TGFBR2* expression was highly expressed by hybrid cells and, as expected, was also more highly expressed by EWS::FLI1-low than EWS::FLI1-high cells (**Fig 3B**). Although *TGFBR2* expression was high in fusion hybrid and low cells, it was detected in only a minority of cells, which may represent true heterogeneity and/or reflect technical limitations of read depth in the scSeq data. To directly measure expression of the receptor protein in EwS cell subpopulations we performed flow cytometry and found that CD73^+^ EwS cells more commonly express TGFBR2 on their surface than CD73^-^ cells (**Fig 3C**).

**Fig 3.**
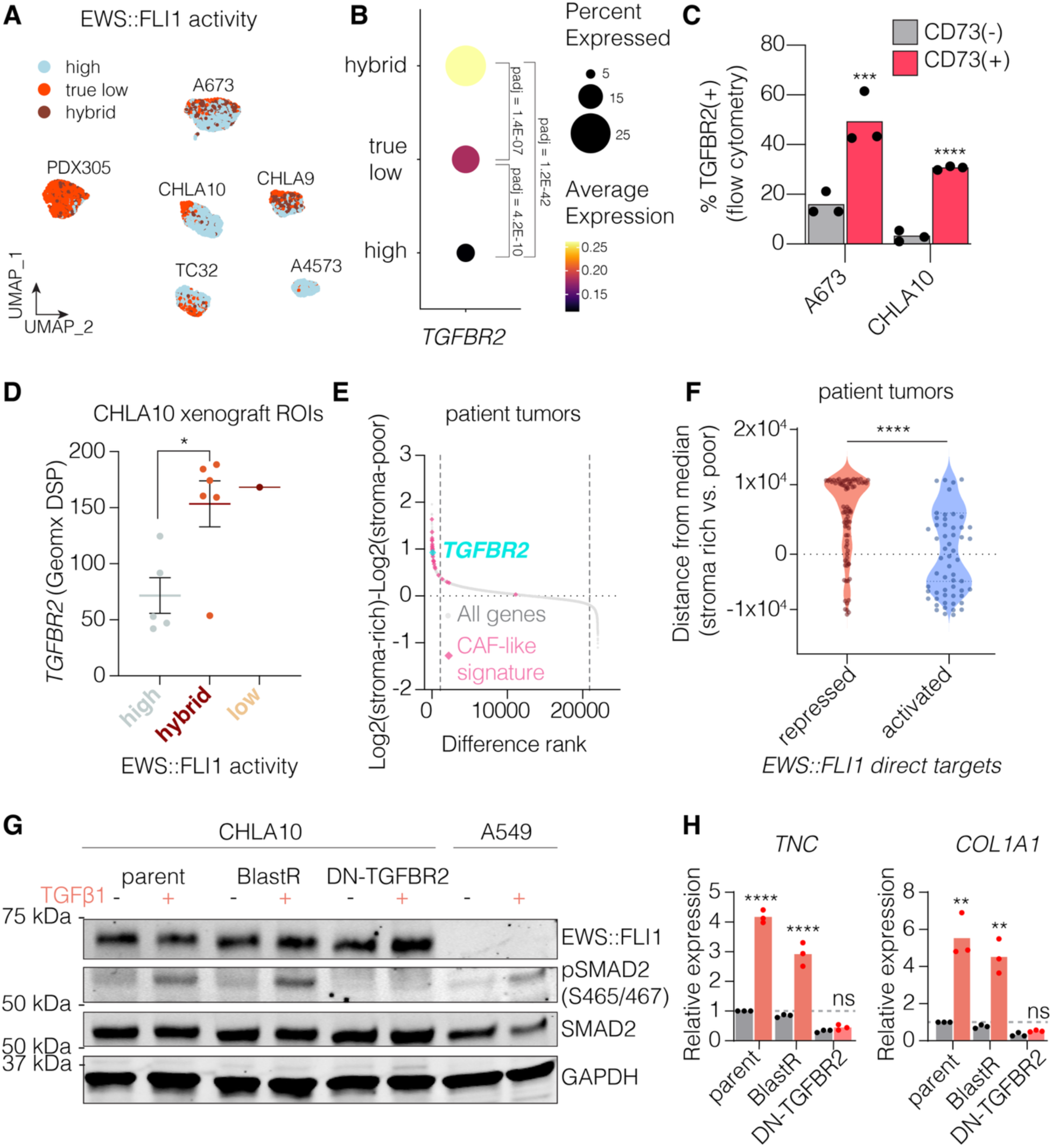
EWS::FLI1-low and -hybrid transcriptional states escape *TGFBR2* repression. (**A**) scSeq data from 6 EwS cell lines with CAF-like subpopulations. Cells scored as EWS::FLI1 “true low” (below median expression of activated/above median expression repressed targets (*17*)) or “hybrid” (above median expression of activated and repressed targets), other cells termed “high”. (**B**) Single cell *TGFBR2* expression grouped by EWS::FLI1 activity state (n=6 cell lines). (**C**) Flow cytometry of TGFBR2 and CD73. n=3 (**D**) *TGFBR2* expression by GeoMx digital spatial profiling of 12 regions of CHLA10 xenografts in NSG mice, previously scored as EWS::FLI1 “low”, “hybrid”, or “high” activity (*10*). (**E**) 40 EwS patient tumors from stroma-poor or stroma-rich (≥30% non-tumor stroma) samples (*36*). Genes ranked by mean stroma-rich/mean stroma-poor sample values. Dashed lines = top/bottom 5% of genes. (**F**) Ranked enrichment in stroma-rich vs. stroma-poor samples in 40 patient tumors (*36*) of EWS::FLI1 activated or repressed target genes (*17*) (0 = not enriched). P-value = Mann-Whitney test. (**G**) Immunoblots of phospho vs. total SMAD2 and EWS::FLI1 in CHLA10 cells transduced with dominant negative TGFBR2 vs. empty vector control -/+ 24 hrs treatment with TGFβ1. Representative of n=2. A549 = positive control for TGFβ induced pSMAD2 and negative control for EWS::FLI1. Loading control = GAPDH. (**H**) RT-qPCR of *TNC* and *COL1A1* in parent, empty vector, or dominant-negative TGFBR2 transduced CHLA10 cells -/+ 24 hrs treatment with TGFβ1. n=3, p-values = unpaired t-tests. ns, *p* > 0.05; *, *p* < 0.05; **, *p* < 0.01; ***, *p* < 0.001; ****, *p* < 0.0001.

Next, we evaluated *TGFBR2* expression in EwS cell subpopulations in vivo. To achieve this we first interrogated transcriptomic data from CHLA10 xenografts that had been subjected to digital spatial profiling (*10*). Data from 12 independent captured regions of interest (ROIs) that were subjected to whole human transcriptome analysis revealed *TGFBR2* expression to be significantly elevated in ROIs that expressed high levels of both EWS::FLI1-activated and EWS::FLI1-repressed genes (**Fig 3D**). Next, we evaluated published gene expression profiling data from pathologically annotated patient tumor biopsies (*36*). Expression of *TGFBR2* was elevated in biopsies that contained abundant ECM/stroma (n=10) compared to biopsies that were ECM/stroma poor (n=30) (**Fig 3E**). Though CAF-like signature (**Fig 3E**) and EWS::FLI1-repressed genes (**Fig 3F**) were upregulated in ECM/stroma-rich biopsies, expression of EWS::FLI1-activated genes did not differ between the two groups (**Fig 3F**). These data from in vivo tumors corroborate in vitro findings that transcriptionally hybrid CAF-like EwS cell subpopulations express higher levels of *TGFBR2* than bulk cell populations.

To assess if TGFBR2 expression is required for TGFβ-induced activation of the CAF-like state, we transduced EwS cell lines with a dominant-negative TGFBR2 construct (DN-TGFBR2) (*37*). DN-TGFBR2 expressing cells failed to activate SMAD2 in response to TGFβ1 (**Fig 3G, Fig S2D**) and induction of ECM gene expression was blocked (**Fig 3H, Fig S2E**). Conversely, expression of EWS::FLI1 was not altered (**Fig 3G, S2D**). These data together demonstrate that *TGFBR2* is expressed by transcriptionally hybrid EwS cells and that TGFβ1-mediated activation of the receptor in these cells induces expression of EWS::FLI1-repressed ECM remodeling genes without significantly altering levels of EWS::FLI1 protein.

### TGFβ induces invasion but not cell cycle arrest in EwS cells

TGFβ induces EMT in malignant epithelial cells and this is most often associated with inhibition of cell proliferation and enhanced invasive potential (*38*). Conversely, in mesenchymal cells TGFβ signaling orchestrates differentiation and can even promote proliferation in some cell types (*39*). To define the phenotypic effects of pathway activation in TGFβ-responsive EwS cells, we first assessed cell proliferation and viability (**Fig 4A-C**). In Incucyte assays, TGFβ1-treated cells reached confluence faster than controls (**Fig 4A**), but this was due to an increase in cell size rather than cell number (**Fig 4B)**. TGFβ1-mediated effects on cell size and confluence were reversed by vactosertib (**Fig 4A,B**). Viability of EwS cells was unaffected by either TGFβ1 or vactosertib (**Fig 4C**).

**Fig 4.**
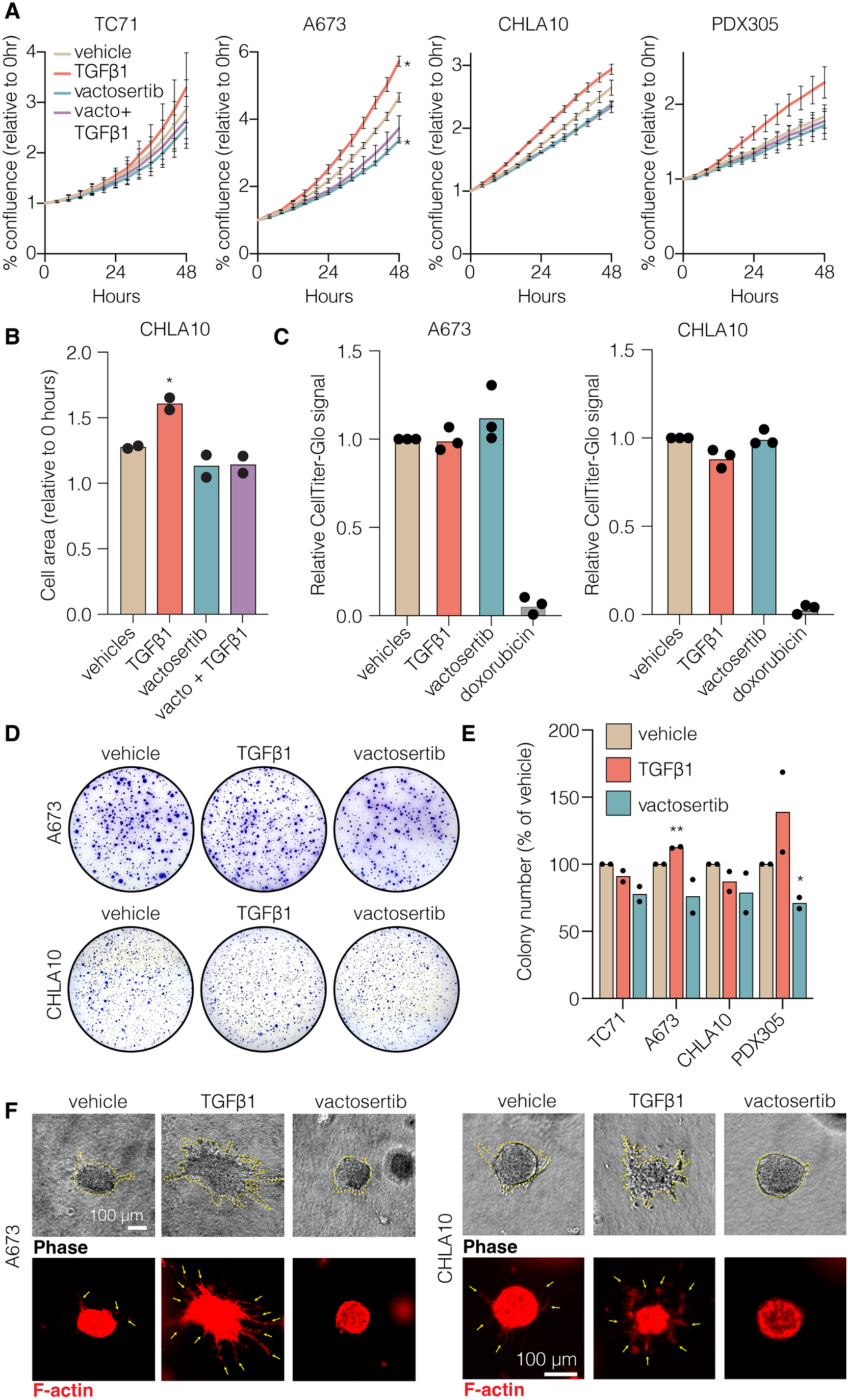
TGFβ induces invasion but not cell cycle arrest in EwS cells. (**A**) Real-time measurements of confluence (Incucyte) for TC71, A673, CHLA10, and PDX305 cells treated with vehicle controls, TGFβ1 (10 ng/mL), vactosertib (1 μM), or TGFβ1 + vactosertib. n=2, error bars = SEM. p-values = unpaired t-tests of confluence at 48hrs relative to 0hrs. (**B**) Measurement of cell area at 48 vs. 0hrs of Incucyte imaging for CHLA10 cells, n=2. Relative to 0hrs. p-value = unpaired t-test. (**C**) Cell viability measurements by CellTiter-Glo after 72 hrs of culture with vehicle control, TGFβ1 (10 ng/mL), vactosertib (1 μM), or doxorubicin (500 nM, positive control for cell death). n=3. (**D**) Soft agar colony forming assays of A673 (day 14) and CHLA10 (day 14) cells treated with vehicle control, TGFβ1 (10 ng/mL), or vactosertib (1 μM). (**E**) Number of colonies formed in soft agar for TC71 (day 14), A673 (day 14), CHLA10 (day 14), and PDX305 cells (day 35), relative to the number of colonies in the vehicle control well. n=2. (**F**) Invasive morphology of spheroids cultured for 4 days in 3D rat-tail collagen I rich gels treated with vehicle controls, TGFβ1 (10 ng/mL), or vactosertib (10 μM). Stained with phalloidin to mark F-actin. Yellow arrows = invasive strands. Representative of n=2. ns, *p* > 0.05; *, *p* < 0.05; **, *p* < 0.01; ***, *p* < 0.001; ****, *p* < 0.0001.

To determine if TGFβ1 altered anchorage-independent growth or invasion we measured colony formation in soft agar (**Fig 4D, E)** and invasive strand formation by spheroids in 3D collagen (**Fig 4F**), respectively. Neither TGFβ1 nor vactosertib altered colony formation of single cells in soft agar (**Fig 4D, E**). Conversely, invasion of EwS spheroids into collagen matrices was strongly augmented by TGFβ1 and reduced by vactosertib (**Fig 4F**). In sum, these data show that activation of TGFβ signaling in EwS cells has no discernible impact on cell proliferation or viability but induces morphologic changes that are associated with increased cell size and enhanced invasive potential.

### TGFβ2 is secreted by CAF-like EwS cells and induces its own expression

Subpopulations of CAF-like EwS cells deposit tumor ECM (*10*) and our data above show that these EWS::FLI1-low and -hybrid cells are more competent to activate TGFβ-dependent transcriptional and phenotypic responses than bulk EWS::FLI1-high cells. Notably, cell lines with high basal expression of the CAF-like signature (CHLA10, PDX305) showed evidence of SMAD2 activation in standard culture conditions prior to the addition of TGFβ1 (**Fig 1G**) leading us to hypothesize that CAF-like EwS cells might secrete TGFβ ligands to activate autocrine signaling. In support of this, flow cytometry evaluation of TGFβ activity using a SMAD binding element (SBE)-GFP reporter revealed significant basal activation of SBE elements in vitro in both CHLA10 and PDX305 cultures in the absence of exogenous ligand (**Fig 5A, Fig S3A**). Single cell transcriptomic data revealed equivalent *TGFB1* expression by EwS cells, irrespective of cell line or transcriptional state (**Fig 5B, Fig S3B).** By contrast, *TGFB2* was almost uniquely expressed by CAF-like cells (**Fig 5B, Fig S3C,D**). ELISA assays detected TGFβ1 protein in the conditioned media of all cell lines, though at levels only slightly higher than unconditioned media, demonstrating that fetal bovine serum serves as the major source of TGFβ1 in culture (**Fig 5C**). Conversely, TGFβ2 was detected in neither fresh nor conditioned media of TC71 or A673 cultures but was abundant in conditioned media of CHLA10 and PDX305 cells (**Fig 5D**). Thus, CAF-like EwS cells express and secrete TGFβ2, creating the potential for autocrine signaling in TGFβ-responsive/TGFBR2*+* cells.

**Fig 5.**
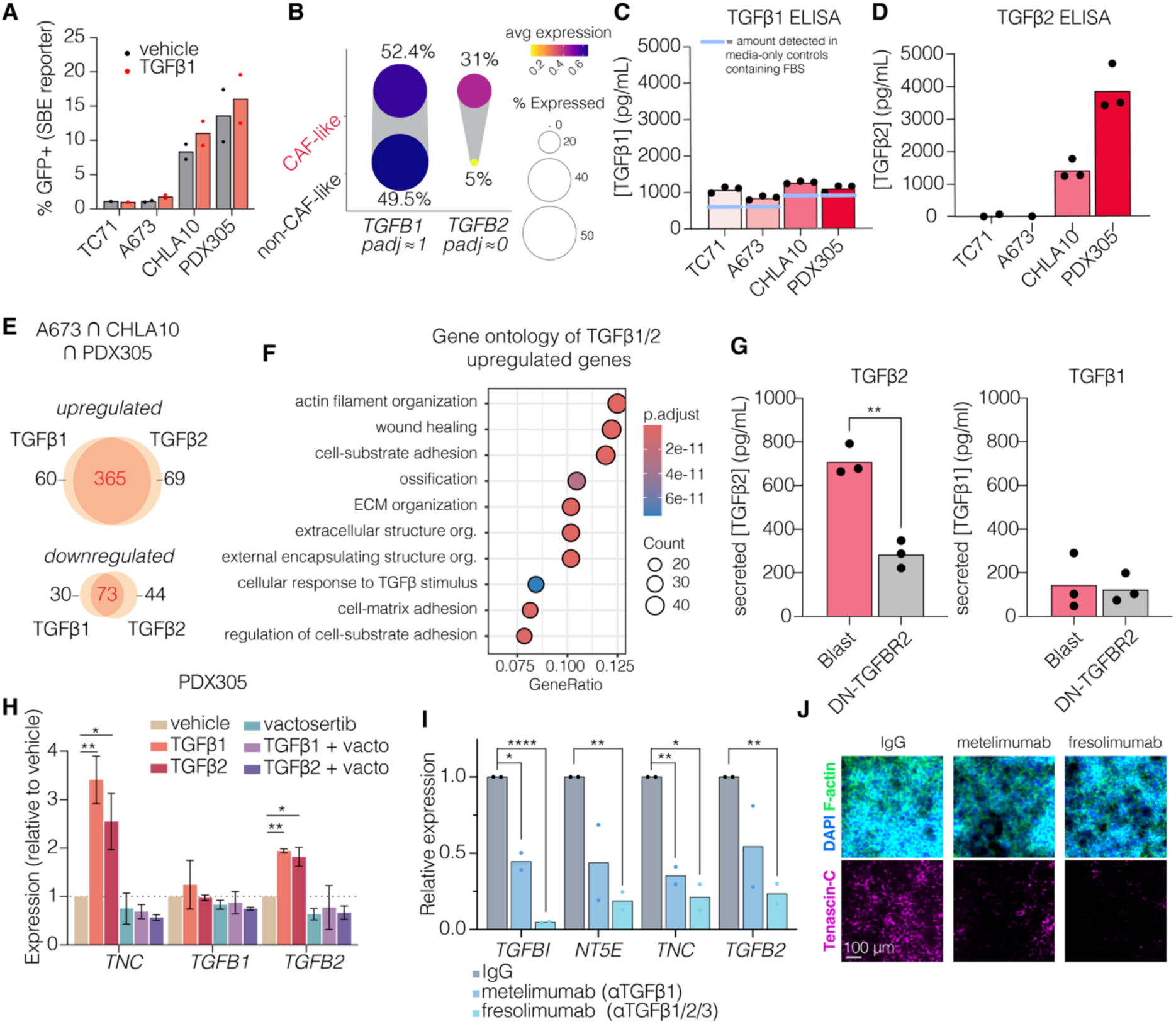
TGFβ2 is secreted by CAF-like EwS cells and induces its own expression. **(A**) % GFP+ after transduction with SBE-GFP SMAD reporter-/+ TGFβ1 (48 hours, n=2). (**B**) *TGFB1* and *TGFB2* expression in CAF-like (top 30% expression CAF-like gene signature) vs. non-CAF-like cells by single cell sequencing (9 EwS lines). (**C**) ELISAs of TGFβ1 in conditioned media (3 days of culture). n=3. Dashed line indicates TGFβ1 level detected in media only controls. (**D**) ELISAs of secreted total TGFβ2 in conditioned media (3 days of culture). n=3. TGFβ2 not detected in media only controls. (**E**) Genes significantly (padj < 0.05) upregulated by TGFβ1 or TGFβ2 in A673, CHLA10, and PDX305 (bulk RNAseq, n=3). (**F**) Gene ontology (GO:BP) of the 365 genes commonly significantly induced (padj<0.05) by TGFβ1 and TGFβ2 in A673, CHLA10, and PDX305 cells. (**G**) ELISAs of secreted total TGFβ2 after 3 days of culture of CHLA10 dominant negative TGFBR2 or empty vector transduced cells. n=3. p-value = unpaired t-test. (**H**) Expression of *TNC*, *TGFB1*, and *TGFB2* in PDX305 cells treated with TGFβ1 (10 ng/mL) -/+ vactosertib, TGFβ2 (10 ng/mL) -/+ vactosertib, or vactosertib alone (1 μM), n=2, p-values = unpaired t-tests. (**I**). Expression of TGFβ induced genes in CHLA10 cells treated with TGFβ ligand blocking antibodies (10 ug/mL, 4 days). (**J**) IF of TNC in CHLA10 cells treated with TGβ ligand blocking antibodies vs. IgG control (10 ug/mL, 4 days). ns, *p* > 0.05; *, *p* < 0.05; **, *p* < 0.01; ***, *p* < 0.001; ****, *p* < 0.0001.

The active cleaved ligand portions of TGFβ1 and TGFβ2 are highly homologous, and in other cell types induce similar gene programs (*40*). Consistent with this, RNAseq profiling of EwS cells showed that the effects of TGFβ2 on EwS cell transcriptomes mirror those of TGFβ1 (**Fig 5E**; **Supplemental Table 3**). Genes induced by both ligands were highly enriched for ECM remodeling, TGFβ pathway response, and wound healing gene ontologies (**Fig 5F**). Like TGFβ1, TGFβ2 did not affect colony formation in soft agar but strongly induced invasion in 3D collagen (**Fig S3E,F**). TGFβ2 also significantly induced expression of the CAF-like gene signature in all cell lines except TC71 (**Fig S3G**). These data support the conclusion that TGFβ1 and TGFβ2 ligands regulate similar transcriptional programs in EwS and that both function to enforce CAF-like phenotypes in TGFβ-responsive EwS cells.

Interestingly, we noted that exposure of CHLA10 and PDX305 cells to either TGFβ1 or TGFβ2 upregulated expression of the *TGFB2* transcript (**Supplementary Tables 1 & 3**). We therefore hypothesized that TGFβ-induced secretion of TGFβ2 by CAF-like EwS cells might activate a positive feedback loop. Consistent with this, interrupting TGFβ signaling in CHLA10 cells via expression of DN-TGFBR2 inhibited cell-autonomous secretion of TGFβ2 (**Fig 5G**). Conversely, TGFβ1 secretion remained low irrespective of DN-TGFBR2 (**Fig 5G**). This self-reinforcing positive feedback was further validated in PDX-derived (**Fig 5H)** and low-passage primary cells (**Fig S3H**) wherein TGFβ ligand-reproducibly upregulated expression of *TGFB2* and *TNC* in these early passage models, and upregulation was completely blocked by vactosertib (**Fig 5H, Fig S3H**).

To test if either or both ligands are required for maintenance of the CAF-like state, we exposed cells to TGFβ blocking antibodies. While the TGFβ1-specific ligand blocking antibody metelimumab (*41*) reduced expression of *TGFBI, NT5E*, *TNC*, and *TGFB2* in CHLA10 cells, expression of all genes was more potently downregulated by fresolimumab, an antibody that blocks all TGFβ ligands (TGFβ1, TGFβ2, and TGFβ3) (*42*) (**Fig 5I**). Fresolimumab also more effectively inhibited TNC protein deposition (**Fig 5J**). Together these findings demonstrate that CAF-like EwS cells produce TGFβ2 which serves to enforce a positive feedback signaling loop in TGFβ-responsive cell subpopulations. Exposing cells to TGFβ blocking antibodies or TGFβ receptor inhibitors interrupts the signaling axis and blunts activation of CAF-like gene signatures and phenotypes.

### Expression of TGFβ2 by EwS cell subpopulations antagonizes fusion-mediated repression of TGFβ signaling

We next investigated the impact of TGFβ2 on expression of EWS::FLI1-modulated genes. Like TGFβ1, TGFβ2 reproducibly induced upregulation of the EWS::FLI1-repressed signature (**Fig 6A**). Effects were again most reproducible and significant in CHLA10 and PDX305 cells where TGFβ2 also inhibited expression of the EWS::FLI1-dependent activation signature (**Fig 6A**). As discussed above, EWS::FLI1 directly represses transcription of *TGFBR2* and our data show that, in TGFβ-responsive EwS cells, both TGFβ1 and TGFβ2 function to inhibit fusion-mediated repression of their receptor (**Fig 6B**). Consistent with the cell autonomous production of TGFβ2 in these lines, vactosertib significantly reduced *TGFBR2* expression from baseline in CHLA10 and PDX305 (**Fig 6B**). To determine if EWS::FLI1 reciprocally regulates expression of TGFβ ligands, we knocked down the fusion in CHLA10 cells and found that *TGFB2* was strongly upregulated concurrently with *TGFBR2* while expression of *TGFB1* was modestly diminished (**Fig 6C**). Upregulation of *TGFB2* and *TGFBR2* in EWS::FLI1-knockdown cells was inhibited by vactosertib (**Fig 6D**) indicating that higher expression of the ligand and receptor in EWS::FLI1-low cells is, in part, dependent on their capacity to activate the TGFβ signaling axis. In support of this, addition of exogenous TGFβ2 to EWS::FLI1-knockdown cells further augmented expression of both *TGFB2* and *TGFBR2* (**Fig 6E**). Importantly, however, while TGFB2-dependent, feedback signaling served to antagonize EWS::FLI1-mediated repression of *TGFB2* and *TGFBR2*, it had no impact on expression of EWS::FLI1-activated genes that are essential for maintenance of tumorigenicity (**Fig 6F**). Moreover, knockdown of EWS::FLI1 in TC71 cells not only led to upregulated expression of both *TGFBR2* and *TGFB2,* it also converted cells from entirely unresponsive to highly TGFβ-responsive cell states (**Fig 6G)**.

**Fig 6.**
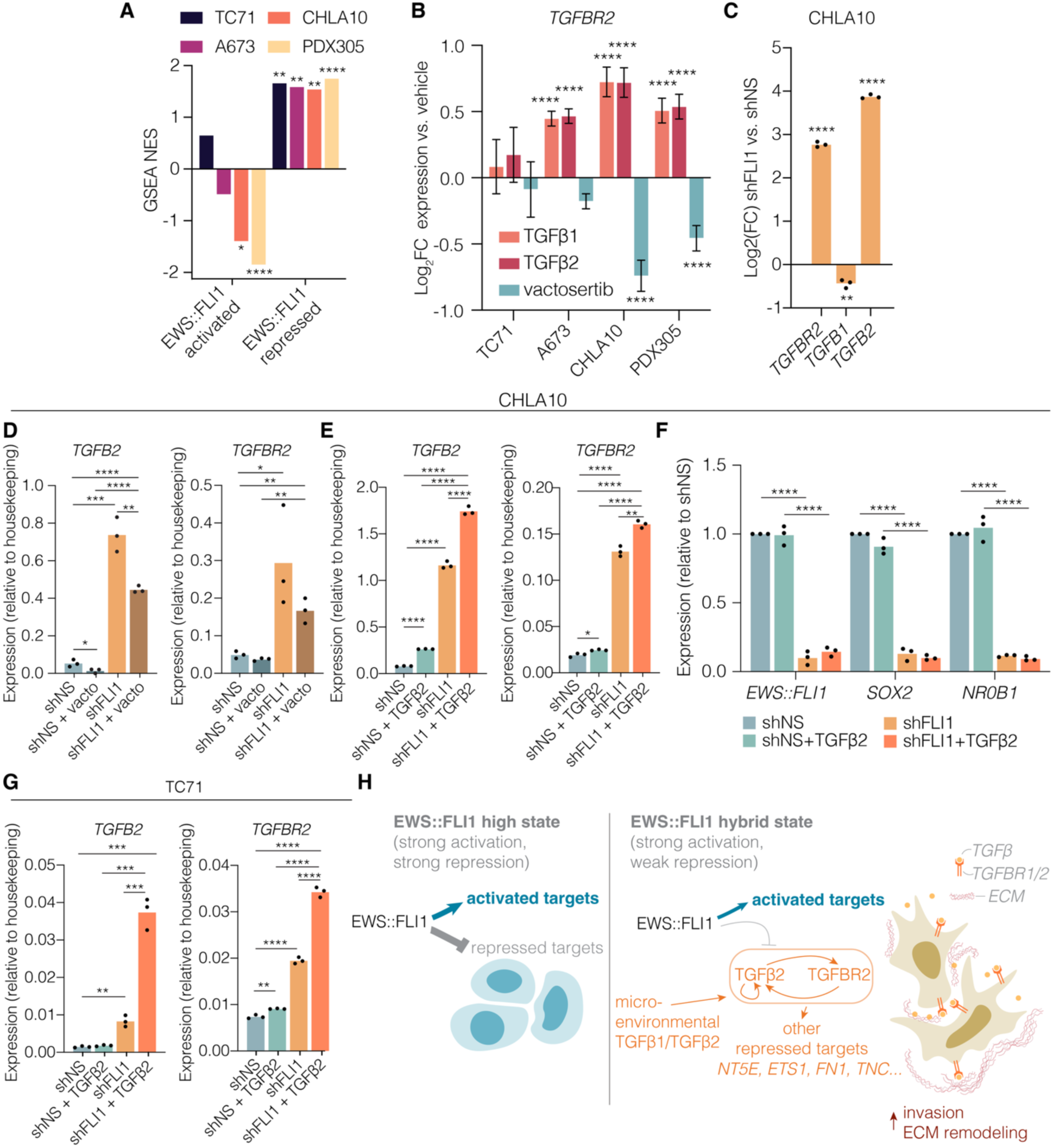
Expression of TGFβ2 by EwS cell subpopulations antagonizes fusion-mediated repression of TGFβ signaling. **(A**) GSEA NES values of EWS::FLI1 target genes (*17*) in TGFβ2 treated cell lines (bulk RNAseq, n=3, p-values = padj from GSEA). (**B**) *TGFBR2* expression (RNA-seq) of cell lines treated with TGFβ1, TGFβ2, or vactosertib (n=3, error bars = standard error, p-values = padj). (**C**) Expression of *TGFBR2*, *TGFB1*, *TGFB2* in shFLI1 vs. shNS CHLA10 cells by RT-qPCR (n=3). (**D**) RT-qPCR of *TGFB2* and *TGFBR2* in CHLA10 cells transduced with shNS or shFLI1 shRNAs (48hrs) then treated with vactosertib (1μM, 24 hrs). n=3. (**E**) RT-qPCR of *TGFB2* and *TGFBR2* in CHLA10 cells transduced with shNS or shFLI1 shRNAs (48hrs) then treated with TGFβ2 (10 ng/mL, 24 hrs). n=3. (**F**) RT-qPCR of *EWS:FLI1* and its direct activated targets *SOX2* and *NR0B1* in CHLA10 cells transduced with shNS or shFLI1 shRNAs (48hrs) then treated with TGFβ2 (10 ng/mL, 24 hrs). Normalized to housekeeping genes then to the average of shNS samples. n=3. P-values = unpaired t-tests. (**G**) RT-qPCR of *TGFB2* and *TGFBR2* in TC71 cells transduced with shNS or shFLI1 shRNAs (48hrs) then treated with TGFβ2 (10 ng/mL, 24 hrs). n=3. (**H**) Model of the positive feedback loop between TGFBR2 activity, TGFβ2 expression, and de-repression of EWS::FLI1 downregulated targets in permissive EWS::FLI1 “low” or EWS::FLI1 “hybrid” states vs. EWS::FLI1 “high” cells. ns, *p* > 0.05; *, *p* < 0.05; **, *p* < 0.01; ***, *p* < 0.001; ****, *p* < 0.0001.

Thus, these data together identify the existence of a positive feedback TGFβ signaling loop in EWS::FLI1-low and EWS::FLI1-hybrid cell subpopulations. Loss of EWS::FLI1-mediated repression of *TGFBR2* sensitizes EwS cells to TGFβ ligands, activating a bidirectional positive feedback loop that induces and sustains expression and secretion of TGFβ2, which in turn antagonizes EWS::FLI1-mediated repression of both *TGFB2* and *TGFBR2*. The phenotypic outcome of amplified TGFβ signaling is creation of invasive and ECM-remodeling EwS cells that maintain expression of pro-tumorigenic EWS::FLI1-activated genes (**Fig 6H**).

## Discussion

Illuminating drivers of transcriptional and phenotypic heterogeneity has the potential to identify biologic programs that are essential for tumor progression. Here we show that TGFβ signaling promotes plasticity of subpopulations of EwS cells, inducing transcriptionally hybrid cells that express pro-metastatic, ECM remodeling genes that are normally repressed by EWS::FLI1. These transcriptionally hybrid cells are more invasive in collagen matrices but proliferation is unaltered, demonstrating that these transcriptionally hybrid cells are also phenotypically hybrid and can both ‘go’ and ‘grow’ (*43*). Our data also reveal that the source of TGFβ ligands can be either stroma- or tumor cell-derived. In particular, we show that CAF-like EwS cells secrete TGFβ2, which then induces cells to transcriptionally upregulate both *TGFB2* and *TGFBR2*, as well as EWS::FLI1-repressed ECM remodeling genes. EWS::FLI1 repressed genes induced by TGFβ include genes that have been previously linked to pro-metastatic phenotypes in the EWS::FLI1 low state. These include *TNC,* which promotes tumor engraftment and metastatic outgrowth (*11*), and migration-related genes *ETS1* (*44*), *ZYX* (*16*), *TEAD3/4,* and other YAP/TAZ dependent genes (*12*). Successful transcriptional activation of TGFβ target genes in both normal development and cancer is highly dependent on cooperation between SMAD proteins and additional transcription factors and epigenetic factors that are active in specific cell contexts (*31, 45*). Whether and how SMAD proteins and other key co-factors participate in antagonizing EWS::FLI1 repressive activity requires further investigation.

A key finding from our studies is that EwS models and patient samples vary widely in their homeostatic proportion of cells that are engaged in these TGFβ-activated and responsive states. A recent spatial transcriptomics assessment of localized and metastatic EwS patient samples likewise found highly spatially variable expression of ECM and EMT genes, including *TNC* and *TGFB1* (*46*). We do not yet know the factors that determine whether individual EwS cells in a cell line or tumor will be TGFβ responsive or non-responsive, but we hypothesize that these homeostatic set points are likely to depend on multiple cell intrinsic and extrinsic parameters that define a cell state equilibrium (*47*). In particular, we speculate that a highly context-dependent interplay between the fusion and other transcriptional regulators in each individual tumor cell in its unique microenvironment explains why TGFβ response differs so widely both within and between EwS cells and patient tumors.

Importantly, our experiments in TGFβ-unresponsive TC71 cells indicate that inhibiting EWS::FLI1 in these cells can convert them to TGFβ-responsive states. Thus, a major determinant of the capacity of an individual EwS cells to engage the TGFβ-signaling loop is its level of fusion expression or activity, which moderates the degree of *TGFBR2* repression. A recent report suggests that even minor losses in EWS::FLI1 more profoundly affect repressed vs. activated genes (*14*), indicating that major downregulation of the fusion to a fully EWS::FLI1-low state may not be required to sensitize cells to TGFβ pathway activation. This is of particular interest in the context of our own findings that EwS cell subpopulations can themselves secrete abundant TGFβ2 protein and that, like *TGFBR2,* expression of the *TGFB2* transcript is repressed by the fusion. Data from published gene expression profiling studies of EWS::FLI1-knockdown cells support our observation that *TGFB2* transcription is subject to EWS::FLI1-mediated repression (*48–50*). In addition, interrogation of published ChIP-Seq data (*51*) reveals intragenic binding of the fusion at the *TGFB2* locus in some cell lines. Further studies are now required to address whether *TGFB2* is a direct or indirect target of EWS::FLI1. Regardless, these findings together have important implications for EwS therapeutics that aim to antagonize EWS::FLI1. Incomplete inhibition of fusion expression or activity may have the unwanted effect of sensitizing EwS cells to activation of a TGFβ2-mediated signaling loop which could, in turn, augment metastatic potential. We propose that, in the absence of complete elimination of the fusion, EWS::FLI1-targeting approaches may benefit from being combined with TGFβ inhibitors or other biologically informed therapeutic combinations (*14*).

TGFβ has not previously been considered a druggable pathway in EwS given fusion-mediated repression of TGFBR2. However, recent work that has highlighted the functional contributions of EWS::FLI1-low cell states to EwS tumor biology has renewed interest in biologic programs that are otherwise downregulated in bulk EWS::FLI1-high cells (*13, 15*). Our group previously showed that activation of canonical Wnt/beta-catenin signaling induces some EwS cells to upregulate secretion of TGFβ2, as well as pro-angiogenic ECM proteins, and that this Wnt-dependent response is mediated, at least in part, by de-repression of *TGFBR2* (*26, 52*). In these prior studies and our current work, we limited our studies to investigate the effects of TGFβ signaling on EwS tumor cells. What now requires deeper investigation is the potential impact of tumor cell-derived TGFβ on non-tumor cells in the TME and where and how stroma-derived TGFβ signals affect EwS cell plasticity. EwS tumors are immunologically cold and generally unresponsive to immune therapies (*53–55*) and TGFβ is a classic negative regulator of the adaptive immune response (*56*). A recent preprint utilizing a humanized immunocompetent orthotopic model of EwS metastasis found that expression of *TGFB1* and *TGFB2* was higher in tumors from humanized immunocompetent models vs. immunocompromised mice (*57*). Encouragingly, a TGFβ-ligand blocking protein (*58*) (which binds both TGFβ1 and TGFβ2) reduced spontaneous metastasis in these humanized mice after radiation treatment. The question remains whether rational incorporation of TGFβ inhibitors into therapeutic regimens might inhibit EwS tumor progression and augment response to immune therapeutics via both tumor cell-intrinsic and TME-dependent mechanisms. Further studies designed to address these critical questions are now warranted.

Taken together, these findings from multiple labs converge to nominate TGFβ as a crucial signal in the EwS TME where it modulates EwS transcriptional plasticity and activation of CAF-like cell states. The data also reveal that CAF-like EwS cells produce TGFβ2 in a feed-forward loop which has implications for creation of metastasis-promoting tumor cell phenotypes and the EwS immune TME landscape. A more detailed mechanistic understanding of the effects of TGFβ on EwS tumor cells and tumor ecosystems will help reveal if and how TGFβ inhibitors could be rationally employed to inhibit metastasis-promoting cell states and augment efficacy of immune and other therapies.

## Materials and Methods

### Cell culture

EwS cell lines A673, TC71, and CHLA10 were obtained from ATCC and COG (https://www.childrensoncologygroup.org/) cell line repositories. PDX305 was generated from an abdominal soft-tissue metastasis from 17-year-old male patient with EwS. PDX305 cells were passaged as a patient-derived xenograft in NSG mice before cell line generation (*10*). A673 and TC71 were maintained in RPMI1640 media (Gibco) supplemented with 10% FBS (Atlas Biologicals) and 2 mmol/L L-glutamine (Life Technologies). CHLA10, PDX305, and PSaRC219 cells were maintained in IMDM media (Fisher) supplemented with 20% FBS, 2 mmol/L L-glutamine, and 1X Insulin–Transferrin–Selenium–Ethanolamine (Gibco). A549 cells (ATCC, lung carcinoma cell line) were cultured in DMEM High Glucose (Gibco) supplemented with 10% FBS. Cells were cultured at 37°C with 5% CO_2_. Cells were all confirmed to be *Mycoplasma*-free and identities subject to short tandem repeat profiling confirmation every 6 months. PSaRC219 low-passage primary patient derived cells (a generous gift from Dr. Bailey, these cells contain a type I EWS::FLI1 fusion and are derived from a lung relapse (*59, 60*)) were cultured on fibronectin-coated plates (human fibronectin: Millipore Sigma 11051407001, coated for 10 minutes at room temperature in PBS at 5.3 μg/mL concentration).

### RT-qPCR

Total RNA was extracted from cells using the RNeasy Mini Kit (Qiagen) and cDNA was generated using the iScript cDNA Synthesis Kit (Bio-Rad) according to the manufacturer’s instructions. qPCR was performed using either TaqMan Fast Universal PCR Master Mix (Applied Biosystems) assays or iTaq Universal SYBR-Green Supermix (Bio-Rad) on a Roche Light-Cycler 480 instrument (Roche Applied Science). Samples were run in technical triplicates and average Ct values were normalized to the geometric mean of two housekeeping genes (*18S* and *EEF1A1* for SYBR Green assays*, 18S* and *B2M* for TaqMan assays). The relative mRNA expression was calculated by the ddCt method when compared to untreated or vehicle controls. Expression relative to housekeeping was used when comparing across cell lines (100*2^-dCt^)). For primer sequences see Supplemental Table 4.

### Incucyte time-lapse imaging assays

5000 cells were plated per well (2000 cells per well for A673 cell line to account for faster doubling rate) in 200 μL complete media in a 96 well plate cultured overnight. The following day media was replaced with 200 μL complete media including treatments and vehicle controls. 10X phase images were collected every 2 hours for 48 hours using an Incucyte (Sartorius). Confluence was measured using the Basic Analyzer (AI Confluence), cell area and cell count were calculated using Adherent Cell-by-Cell analysis optimized for each cell line.

### CellTiter-Glo assessment of cell viability

5000 viable single cells were plated per well of opaque white walled 96 well plates (when A673 cells were used 2000 cells were plated per well to account for faster doubling rates) and cultured overnight. Wells were then treated for 72 hours in 200 μL total volume complete media -/+ appropriate treatments. 80 μL of CellTiter-Glo was added per well and incubated for at least 10 minutes at room temperature. Total luminescence was measured (1000 ms integration time) on a SpectraMax iD5 (Molecular Devices) plate reader.

### Colony formation in soft agar

5000 single cells were plated in 0.45% UltraPure low melting point agarose (Invitrogen 16520-050) at 37°C on top of a solidified bottom layer of 1mL 5% agarose in 6 well plates. Gels were solidified by incubation at 4°C for 10 minutes then supplemented with 1mL complete media -/+ treatments. Media was changed twice per week until visible colonies had formed (14 days for TC71, A673, and CHLA10; 35 days for the slower growing PDX305 cell line). Colonies were stained with 300 μL of 0.1% crystal violet solution in 20% methanol for 20 minutes at room temperature, then washed 5 times for 5 minutes and once overnight with distilled H_2_O. Well images were thresholded in FIJI and colony number and area were measured using the Analyze Particles function.

### 3D rat tail collagen invasion assays

Spheroids were formed by plating EwS single cells at a density of 50,000 viable cells/mL in 4mL complete media in ultra-low-attachment 6 well plates overnight (Thermo Fisher Scientific 07-200-601). Debris and single cells were removed by centrifuging samples for 10 seconds at 300g the following day. Rat tail collagen (Gibco) was prepared at 1.8 to 2 mg/mL in 0.08 N acetic acid, supplemented with 10X DMEM (final concentration 1X), neutralized with 0.5 mol/L NaOH and allowed to polymerize on ice for 45 to 60 minutes until fibers formed (*61*). Spheroids and collagen gels were mixed and plated as 80 μL droplets and allowed to polymerize for 30 minutes at 37°C, at which point 1 mL of complete IMDM media (supplemented with 20% FBS, 1X L-Glutamine, 1X ITS-X) was added. 20 μL underlays of polymerized rat tail collagen were first applied to the bottom of a tissue culture– treated 24wp plate to prevent spheroids in the overlying collagen dome from attaching to or spreading along the plate surface. Spheroids were cultured 4 days then assessed by phase contrast microscopy and fixed for 15 minutes with 4% PFA in PBS. Invasion was scored as protrusive cells or multicellular strands extending from the spheroid border. Spheroids were stained for at least 1 hour at room temperature with DAPI (2.5 μg/mL) and Alexa fluorophore conjugated phalloidin (Thermo Fisher, 400X) to localize nuclei and F-actin, respectively.

### Renal subcapsular injection xenografts

NSG male mice (Jackson Laboratory, 005557) were anesthetized using 2% isoflurane in O2 delivered at 1.5 L/min and their flank shaved. Animal was secured to its side on the heated platform of the ultrasound VEVO 3100 (VisualSonics Inc) and the MX550D transducer positioned to get an image of the kidney. Using the image from the ultrasound, a 27g needle was gently inserted through the skin and back muscle into the sub-renal capsule space and 5000 A673-GFP-Luciferase cells in 15 μL PBS were mixed with 15 μL Matrigel on ice then injected using an insulin syringe. After 1min, the needle was slowly removed and the animal recovered in a warmed cage. The size of the tumor was monitored weekly by ultrasound and the animals were humanely sacrificed when the tumor reached approximately half the size of the kidney (3 weeks post-injection).

### Immunofluorescence

#### Cells

For immunofluorescence, cells were fixed with 4% PFA in PBS for 10 minutes then washed with PBS. Cells were blocked with 0.2% BSA in PBS for 1 hour at room temperature. Primary antibodies were incubated for 1 hour at room temperature or overnight at 4°C in 0.2% BSA (Supplemental Table 4 Key Reagents Table). Alexa-fluor–conjugated secondary antibodies were incubated for 1 hour at room temperature in 0.2% BSA. Signal was compared with matched species-specific IgG controls incubated at the same concentrations. For F-actin staining Alexa Fluor conjugated phalloidin was used (Thermo Fisher, 400X). DAPI was included during secondary antibody incubations to mark nuclei at 2.5 μg/mL. Coverslips were mounted with ProLong Gold mounting media and allowed to cure overnight at room temperature in the dark. Slides were imaged on a Leica DMi8 Thunder Imager widefield deconvolution microscope.

#### Tissue

Formalin-fixed, paraffin-embedded (FFPE) tumor samples from xenografts or primary patient samples were deparaffinized, then antigen retrieval was carried out using Diva Decloaker in a Cuisinart CPC-900 (high pressure, 5 minutes) and slides were allowed to cool to room temperature. Slides were blocked with 0.2% BSA for 1 hour at room temperature. Primary and secondary antibodies (Supplemental Table 4 Key Reagents Table) were incubated in 0.2% BSA overnight at 4°C. DAPI was included during secondary antibody incubations to mark nuclei at 2.5 μg/mL. Coverslips were mounted with Prolong Gold and allowed to cure overnight at room temperature in the dark, then slides were stored at 4°C until imaging. Slides were imaged on a Leica DMi8 Thunder Imager at 20X magnification.

### Immunofluorescence quantification

QuPath (version 0.5.1) was used for quantification of pSMAD2 S465/467 expressing cells in a previously generated EwS patient TMA (*10*). A 20X slide scan fluorescent image of the entire TMA was separated into a TMA grid and missing or damaged cores were excluded. Cell detection was performed using the DAPI channel (2.5 micron cell expansion). Measurements for each cell and channel were exported as .csv files and converted to .fcs files using the write.FCS function of the flowCore package in R. Cells were gated on cytoplasmic/membrane CD99 (anti-mouse-Alexa-555) signal and nuclear pSMAD2 (anti-rabbit-Alexa-647) signal in FlowJo 10.8.2.

### Immunoblots

Cells were lysed with RIPA buffer (Fisher Scientific) supplemented with protease and phosphatase inhibitors (Sigma). Western blots were performed using the Bio-Rad Mini-PROTEAN Tetra System. Following transfer, nitrocellulose membranes were blocked in Odyssey Blocking Buffer (LI-COR) for 1 hour. Membranes were washed and incubated rocking overnight at 4°C with primary antibodies (see Supplemental Table 4). Membranes were then washed four times in TBST for 5 minutes each and incubated with a secondary antibody (LI-COR IRDye 700CW or 800CW; 1:10,000) for 1 hour. After 4 additional 5-minute TBST washes membranes were scanned on a LI-COR Odyssey scanner.

### Lentivirus production and transductions

Lentiviruses were produced using a high-throughput, second-generation lentiviral system based on the Daedalus mammalian protein expression system (*62*), adapted for a custom robotics platform. Suspension-adapted HEK 293T cells (2 × 10⁶ cells per mL, 1 mL per well) were seeded into 2 mL round-bottom 96-well blocks (Axygen, P-DW-20-C-S). For each well, 0.87 μg of transfer plasmid, 0.42 μg psPax2 (Addgene #12260), and 0.21 μg pMD2.G (Addgene #12259) were combined with 21 μL polyethyleneimine (PEI, 1 μg/μL) in FreeStyle™ 293 Expression Medium (ThermoFisher, Cat. #12338018) using a Hamilton Starlet liquid handler. The transfer plasmids utilized in this study included pLenti CMV Blast DEST (Addgene #17451), pLenti CMV Blast DN-TGFBR2-HA (Addgene #130888), pLKO.1 shFLI1 (Sigma TRCN0000005322), and pLKO.1 non-silencing control (Sigma SCH002). Following transfection, the blocks were incubated in a Cytomat incubator at 37°C, 5% CO₂, and 8% humidity with agitation at 1,000 RPM. Twenty-four hours after transfection, 0.6 μg of valproic acid was added to each well. On day four, supernatants were collected by centrifugation at 500 × g for 10 minutes and then pooled into 24-well deep-well plates (Whatman, 7701-5102). SBE/SMAD-GFP Puro reporter lentivirus was purchased (LipExoGen LTV-0013-1S, 5 TU added per viable cell for transduction). For transductions, 150,000 viable cells were plated in a 6 well plate. The following day complete media containing 8 μg/mL polybrene was added. 150 μL of unconcentrated lentiviral supernatant was added to each well overnight. For puromycin resistant constructs 48 hours later puromycin was included at 0.5 μg/mL to the culture media for at least 5 days of selection, for BlastR constructs blasticidin was included at 8 μg/mL for at least 10 days.

### Flow cytometry and FACS

EwS cells were trypsinized, washed with PBS, resuspended in 2% FBS in PBS, and stained with fluorophore-conjugated antibodies (Supplemental Table 4 Key Reagents Table) for 30 minutes on ice in the dark. Fluorophore-conjugated isotype controls and unstained controls were used for each experiment. Flow cytometry was performed on a BD Accuri C6 (10,000 events), with single-cell identification by forward scatter and side scatter. For FACS, CD73-PE negative vs. positive cells were sorted on a BD FACSAria II. Analysis was conducted in FCS Express (De Novo Software). Isotype controls were used for conjugated primary antibodies and untransduced (non-fluorescent) cells were used as negative controls for SBE-GFP reporter lines.

### ELISAs

To prepare conditioned media for ELISAs, EwS cells were cultured for 3 days in 6 well plates. Cells were pelleted and supernatant was collected. To activate latent TGFβ ligands 20 μL of 1M HCl was added at room temperature for 10 minutes to each 100 μL of conditioned media, then quenched with 20 μL 1M NaOH/400 mM HEPES (final estimations of ligand concentration in CM accounted for this 1.4x dilution factor). CM was diluted 2-fold in Diluent B, manufacturer instructions were followed (ThermoFisher EHTGFB2, BNS249-4). Cell counts were taken on day 3 to confirm similar viability and total cell count per well across conditions.

### Bulk RNA-seq

Poly(A)-capture RNA-seq was performed on RNA (purified using Qiagen RNeasy kit) from TC71, A673, CHLA10, and PDX305 cells treated with TGFβ1 (10 ng/mL), TGFβ2 (10 ng/mL), vactosertib (1uM), or vehicle controls for 24hrs. Paired end 150 bp sequencing was performed on a NovaSeq X Plus. Library preparation and sequencing was performed by Novogene. Trimgalore 0.6.7, FASTQC 0.11.9, STAR Align 2.7.10a, Picard markduplicates 3.0.0, and Samtools 1.17 were used for FASTQ quality control, trimming, and alignment to reference genome GrCh38. Counts and differential expression analysis were calculated using DESeq2 (v1.44.0). GSEA and gene ontology of GO:BP terms were analyzed using clusterProfiler (version 4.12.6). Venn diagrams were created using Vennerable (version 3.1.0.9000).

### Public tumor transcriptomic datasets

Single cell sequencing of 5 EwS patient derived xenograft subcutaneous tumors in NSG mice (*9*) were obtained from GSE130025. Nanostring GeoMx digital spatial profiling data of CHLA10 xenografts generated by subcutaneous or tail vein injection (GSE226562) and single cell sequencing of 9 EwS cell line (GSE236284) were re-analyzed here (*10*). A subset of 40 EWS::ETS fusion positive patient tumor samples subjected to Affymetrix Human Exon 1.0 ST array expression analysis from GSE63157 were used based on pathological scoring as stroma rich (“invading”, ≥ 30% non-tumor stroma, n=10) or stroma-poor (“core”, n=30) based on H&E staining (*36*).

### Statistical Analysis

Statistical tests were conducted in GraphPad Prism 10. Unpaired *t*-tests were used unless otherwise specified. Error bars represent standard deviation of biological replicates unless otherwise specified. ns, *p* > 0.05; *, *p* < 0.05; **, *p* < 0.01; ***, *p* < 0.001; ****, *p* < 0.0001.

### Study approval

All experimental procedures adhere to the *Guide for the Care and Use of Laboratory Animals* and were approved by Seattle Children’s Research Institute IACUC. The institute is fully AAALAC-accredited and has a Public Health Service approved Animal Welfare Assurance.

## Data and code availability

Sequencing data generated by this study will be available in the NCBI Gene Expression Omnibus database upon publication. Analyses in R will be available upon publication at github.com/LawlorLab.

## Acknowledgments

We thank the patients who generously donated their tissues. We thank Seattle Children’s Research Institute Bioinformatics and Research Scientific Computing, the Microscopy and Histopathology CoLab, and the Flow Cytometry Core.

## Author Contributions

Conceptualization: EDW, ERL

Methodology & Resources: SCC, JSP, JMO, KMB

Formal analysis: EDW, JRV

Investigation: EDW, JCH, AAA, SIW, AM, PAL, NMG

Visualization: EDW

Supervision: ERL

Writing—original draft: EDW

Writing—review & editing: EDW, ERL.

## Data and Materials Availability

All data are available in the main text or the supplementary materials. RNA sequencing data to be submitted to NCBI GEO upon publication, and R code used to generate analyses and figures to be deposited to github.com/LawlorLab.

## SUPPLEMENTAL TABLES

**Supplemental Table 1.** Bulk RNAseq differential expression analysis of TC71, A673, CHLA10, or PDX305 cells treated with TGFβ1, vactosertib, or vehicle control.

**Supplemental Table 2.** Clinical details of patient tumor microarray samples subjected to CD99 and pSMAD2 immunofluorescence.

**Supplemental Table 3.** Bulk RNAseq differential expression analysis of TC71, A673, CHLA10, or PDX305 cells treated with TGFβ2 or vehicle control.

**Supplemental Table 4**. Key reagents table.

