## Supplemental_Figures_S1-S3 for "Autocrine TGFβ2 enforces a transcriptionally hybrid cell state in Ewing sarcoma"

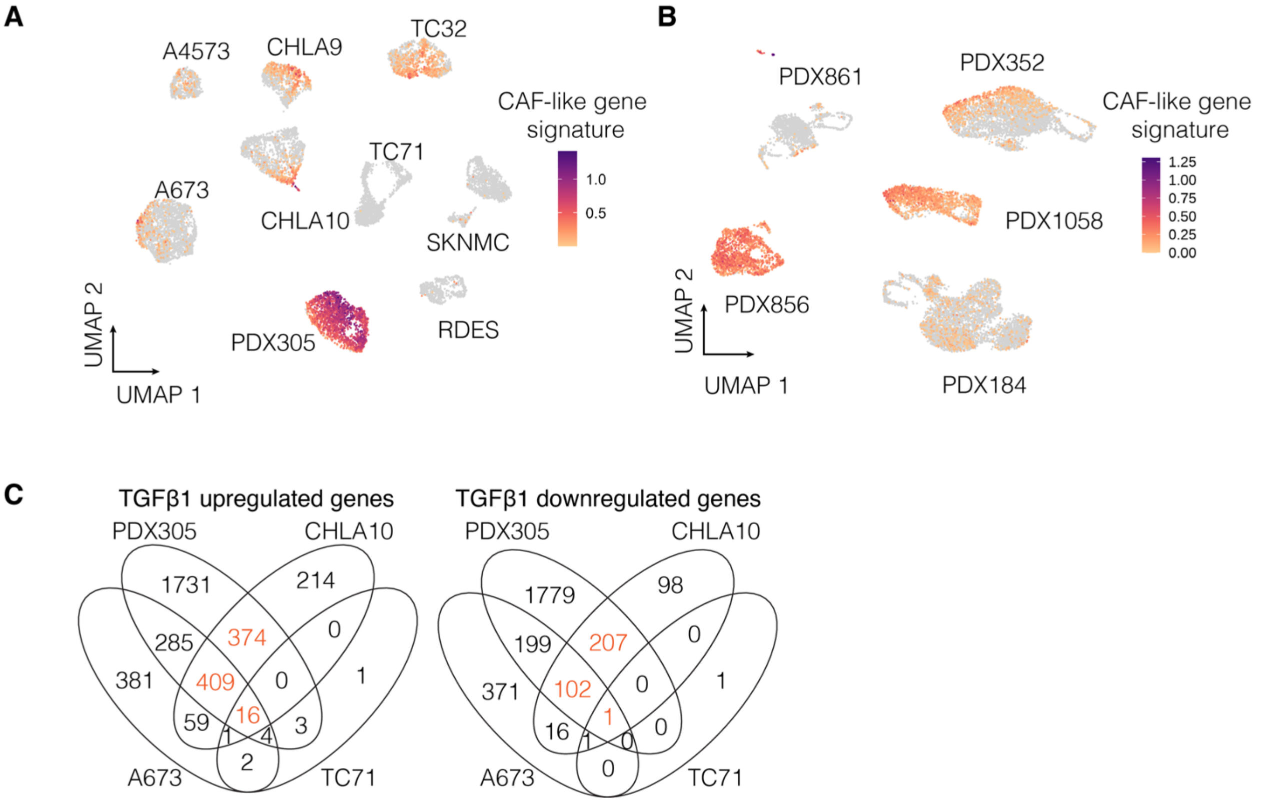


**Figure S1. Related to Figure 1.** (A) UMAP of single cell sequencing data for 9 Ewing sarcoma cell lines showing expression of the 28-gene *NT5E*-associated CAF-like signature (10). (B) UMAP of single cell sequencing data for 5 Ewing sarcoma PDXs in NSG mice (9) showing expression of the 28-gene *NT5E*-associated CAF-like signature from. (C) Overlap of genes significantly up or downregulated (padj<0.05) by 24 hrs TGFβ1 vs. vehicle treatment (bulk RNAseq, n=3) of TC71, A673, CHLA10, or PDX305 cell lines.

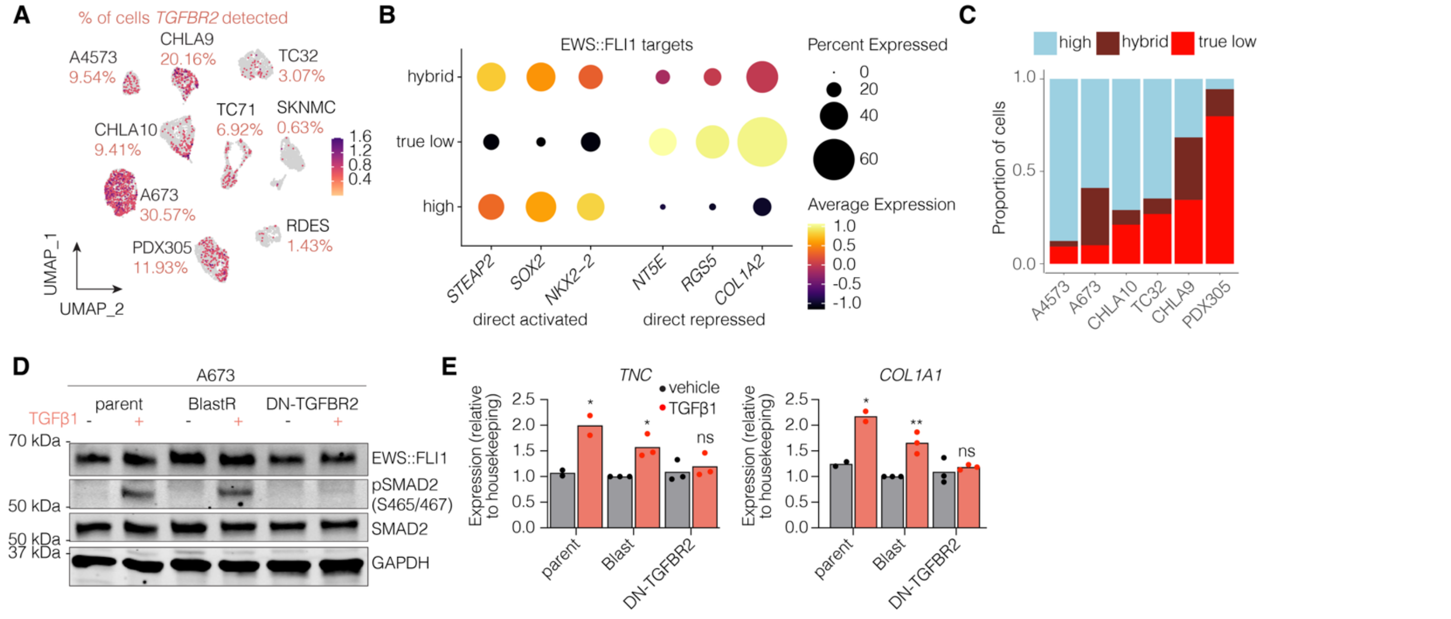


**Figure S2. Related to Figure 3.** (A) Expression of *TGFBR2* (% of cells with *TGFBR2*>0 reported for each line in red) by single cell sequencing of 9 EwS cell lines. (B) Expression of individual EWS::FLI1 activated or repressed targets across EWS::FLI1 activity states for 6 cell lines (A4573, A673, CHLA9, CHLA10, TC32, PDX305). (C) Proportion of cells in different EWS::FLI1 activity states ("high", "hybrid", or "true-low") for 6 EwS cell lines. (D) Immunoblots of phospho vs. total SMAD2 and EWS::FLI1 in A673 cells including untransduced parental control or cells transduced with a dominant negative (kinase dead) TGFBR2 construct to block TGFβ pathway activation vs. an empty vector control -/+ 24 hrs treatment with TGFβ1. Representative of n=2. Loading control = GAPDH. (E) RT-qPCR of *TNC* and *COL1A1* in parent, empty vector, or dominant-negative TGFBR2 transduced A673 cells -/+ 24 hrs treatment with TGFβ1. n=3. Unpaired t-tests. ns, $p > 0.05$; *, $p < 0.05$; **, $p < 0.01$; ***, $p < 0.001$; ****, $p < 0.0001$.

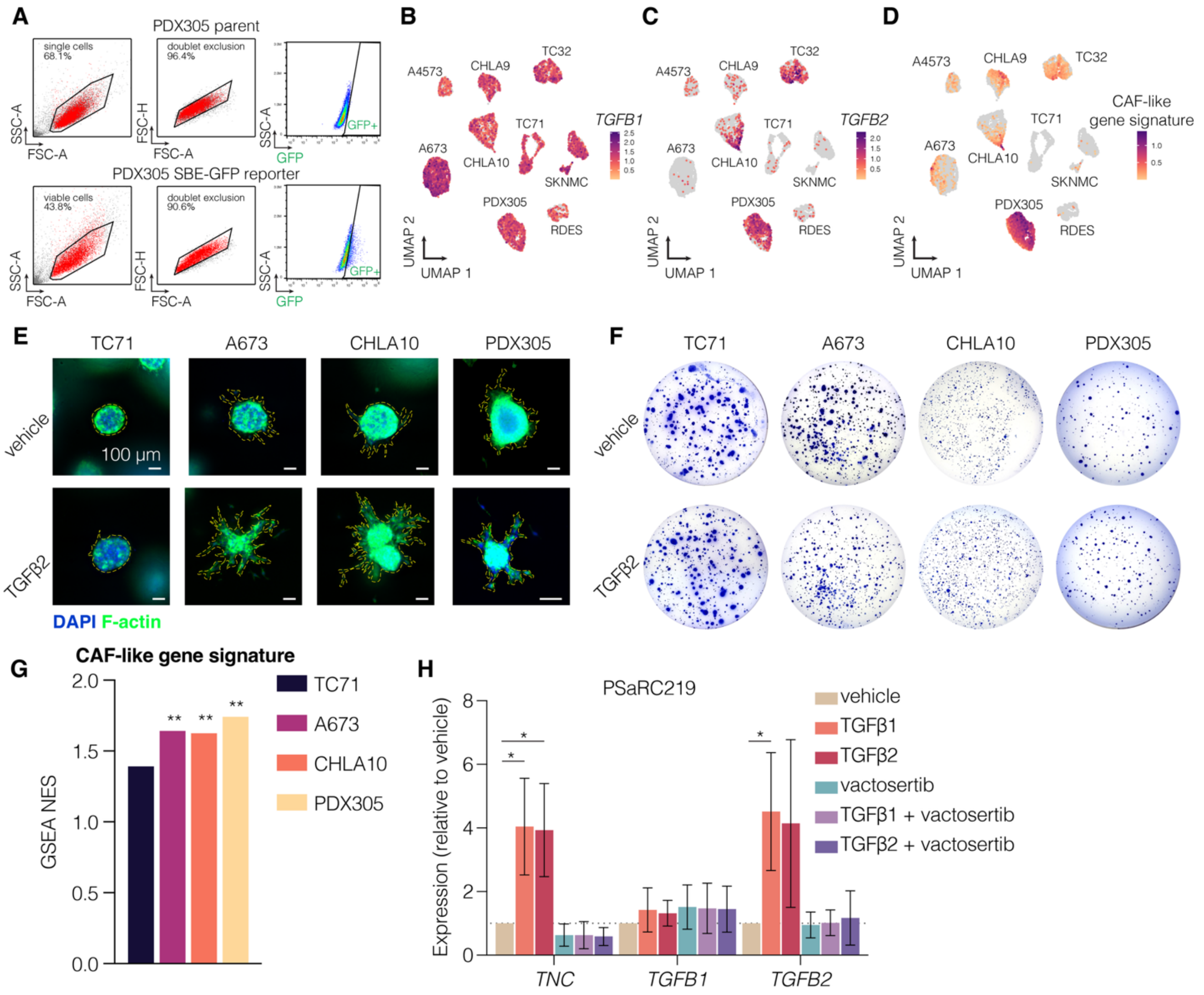


**Figure S3. Related to Figure 5.** (A). Gating strategy for detecting GFP+ cells transduced with the SBE-GFP SMAD activity reporter. Untransduced PDX305 (top) vs. SBE-GFP transduced PDX305 (bottom). Expression of (B) *TGFB1*, (C) *TGFB2*, or (D) the 28-gene CAF-like signature by single cell sequencing of 9 EwS cell lines. (E) Invasive morphology of spheroids cultured for 4 days in 3D collagen gels treated with TGFβ2 (10 ng/mL) or vehicle. Representative of n=2. (F) Soft agar colony forming assays of TC71 (day 14), A673 (day 14), CHLA10 (day 14), and PDX305 cells (day 35) treated with vehicle control or TGFβ2 (10 ng/mL). (G) GSEA NES scores of the CAF-like signature in cells treated with TGFβ2 vs. vehicle. p-values = padj. (H) Expression of *TNC*, *TGFB1*, and *TGFB2* in low passage number primary patient cell line PSaRC219 treated with vehicle control, TGFβ1 (10 ng/mL) -/+ vactosertib, TGFβ2 (10 ng/mL) -/+ vactosertib, or vactosertib alone (1 µM), n=3. p-values = unpaired t-tests. ns, $p > 0.05$; *, $p < 0.05$; **, $p < 0.01$; ***, $p < 0.001$; ****, $p < 0.0001$.
